# *RNA Binding Motif Protein 48* is required for U12 splicing and maize endosperm differentiation

**DOI:** 10.1101/341917

**Authors:** Fang Bai, Jacob Corll, Donya N. Shodja, Ruth Davenport, Guanqiao Feng, Janaki Mudunkothge, Christian J. Brigolin, Federico Martin, Gertraud Spielbauer, Chi-Wah Tseung, Amy E. Siebert, W. Brad Barbazuk, Shailesh Lal, A. Mark Settles

## Abstract

The last eukaryotic common ancestor had two classes of introns that are still found in most eukaryotic lineages. Common U2-type and rare U12-type introns are spliced by the major and minor spliceosomes, respectively. Relatively few splicing factors have been shown to be specific to the minor spliceosome. We found that the maize RNA Binding Motif Protein48 (RBM48) is a U12 splicing factor that functions to promote cell differentiation and repress cell proliferation. RBM48 is coselected with the U12 splicing factor, ZRSR2/RGH3. Protein-protein interactions between RBM48, RGH3, and U2 Auxiliary Factor (U2AF) subunits suggest major and minor spliceosome factors may form complexes during intron recognition. Human RBM48 interacts with ARMC7. Maize RBM48 and ARMC7 have a conserved protein-protein interaction. These data predict that RBM48 is likely to function in U12 splicing throughout eukaryotes and that U12 splicing promotes endosperm cell differentiation in maize.

## Introduction

Eukaryotic protein-coding genes generally are interrupted by introns, and splicing of pre-mRNA is fundamental for protein expression. It has been proposed that eukaryotic introns originate from bacterial, self-splicing Group II introns (Burge et al., 1998; Roy and Irimia, 2009). However, eukaryotic introns are not self-splicing, and dynamic RNA-protein complexes, known as spliceosomes, direct accurate recognition of splice sites (Simpson and Filipowicz, 1996; Brown, 1998; Lorkovic et al., 2000; Ru et al., 2008; Rappsilber et al., 2002). The last eukaryotic common ancestor had two classes of introns that were spliced by different spliceosome complexes (Irimia and Roy, 2014). The vast majority of introns, termed U2-type introns, are spliced by the major spliceosome (Lee and Rio, 2015). There are also rare, U12-type introns, which are spliced by the minor spliceosome (Will and Luhrmann, 2005). U12-type introns have a higher degree of sequence conservation at the 5’ splice site and branch point sequence (BPS) relative to U2-type introns (Turunen et al., 2013).

The minor spliceosome is at a low concentration in the cell, and splicing of U12-type introns is significantly slower than U2-type introns both *in vitro* and *in vivo* (Tarn and Steitz, 1995; Patel et al., 2002; Montzka and Steitz, 1988). Consequently, splicing of U12-type introns can limit the abundance of mature transcripts from minor intron-containing genes (MIGs) (Konig et al., 2007; Patel et al., 2002; Niemela et al., 2014; Niemela and Frilander, 2014; Younis et al., 2013). In human cells, U12 splicing efficiency is responsive to a stress-activated signal transduction pathway (Younis et al., 2013). These data suggest that U12 splicing efficiency could be a deeply conserved regulatory mechanism. However, evolutionary evidence indicates the minor spliceosome is dispensable in many eukaryotes. The minor spliceosome has been lost in multiple independent lineages including model organisms such as *C*. *elegans*, *S. cerevisiae, D. discoideum,* and *C. reinhardtii* (Dávila et al., 2008). Among metazoan species with U12 splicing, the number of MIGs varies drastically, ranging from 758 in humans to only 19 in *D. melanogaster* (Alioto, 2006). Loss of U12-type introns is the most common mechanism for reduction of MIGs within a genome (Burge et al., 1998).

Despite species variation for the presence and number of MIGs, U12 splicing appears to have roles in cell proliferation, cell differentiation, and development across divergent eukaryotes. A striking example is revealed by mutations of ZRSR2 orthologs in human and maize. Somatic mutations in human *ZRSR2* primarily affect U12 splicing and cause myelodysplastic syndrome, a disease that reduces differentiation of mature blood cell types (Madan et al., 2015). The *rgh3* mutation of the maize ZRSR2 ortholog has a conserved U12 splicing function and inhibits differentiation of endosperm cells (Fouquet et al., 2011; Gault et al., 2017). Other U12 splicing mutants in humans and mouse models demonstrate roles in hematopoiesis, muscle strength, as well as bone and neural development (Edery et al., 2011; He et al., 2011; Horiuchi et al., 2018). A mutation in the zebrafish RNPC3 locus disrupts digestive organ development (Markmiller et al., 2014). Arabidopsis knockdowns of minor spliceosome proteins show defects in leaf development and flowering time (Kim et al., 2010; Jung and Kang, 2014; Xu et al., 2016). The different pleiotropic phenotypes of U12 splicing factors have made it difficult to identify unifying developmental functions for MIGs.

Moreover, there are relatively few splicing factors that have been identified as specific to the minor spliceosome. Biochemical characterization of human small nuclear ribonucleoproteins (snRNPs) showed that most minor spliceosome proteins are shared with the major spliceosome with only seven novel proteins uniquely associated with the minor spliceosome: PDCD7, RNPC3, SNRNP25, SNRNP35, SNRNP48, ZCRB1, and ZMAT5 (Schneider et al., 2002; Will et al., 2004). Genetic analysis of human ZRSR2, human FUS, mouse SMN1, and maize RGH3 indicates that these factors predominantly affect U12-type intron splicing even though there is substantial evidence for the proteins to participate in other RNA metabolic processes (Gault et al., 2017; Madanieh et al., 2015; Reber et al., 2016). Here we identify the maize RNA Binding Motif Protein (RBM48) as a minor spliceosome factor which functions to promote cell differentiation and repress cell proliferation. RBM48 is conserved in organisms that retain the minor spliceosome predicting that this protein will have a function in U12 splicing throughout eukaryotes. Protein-protein interactions and co-localization between RBM48, RGH3, and U2 Auxiliary Factor (U2AF) subunits suggests major and minor spliceosome factors may form complexes as part of recognizing introns.

## Results

### Loss of RBM48 causes a *rough endosperm* (*rgh*) defective kernel phenotype

The *rbm48-umu1* reference allele was identified from a collection of 144 UniformMu *rgh* mutants (Fouquet et al., 2011; McCarty et al., 2005). The *rgh* locus was mapped to the long arm of chromosome 4 with bulked segregant analysis (Liu et al., 2010), and compared to *Robertson’s Mutator* transposon flanking sequence tags from the *rgh* mutant line (Supplementary Figure 1). The *rbm48-umu1* insertion in GRMZM2G163247 was the only novel insertion in this line that co-segregated with the *rgh* phenotype (Supplementary Figure 1). The *rbm48-umu2* allele was obtained from the Maize Genetics Co-operative Stock Center (McCarty et al., 2013). Self-pollination of *rbm48-umu2* heterozygotes also segregate for an *rgh* kernel phenotype (Supplementary Figure 1). Both alleles show similar defective kernel phenotypes and segregate at ratios consistent with a recessive mutant (Figure 1, Supplementary Table 1). The *rbm48-umu1* allele transmits fully through both male and female gametes but does not give a seed phenotype when crossed to normal inbred lines (Supplementary Table 2). Crosses of the two alleles failed to complement the *rgh* mutant phenotype indicating that *rbm48* mutations disrupt maize seed development (Supplementary Figure 2).

**Figure 1.**
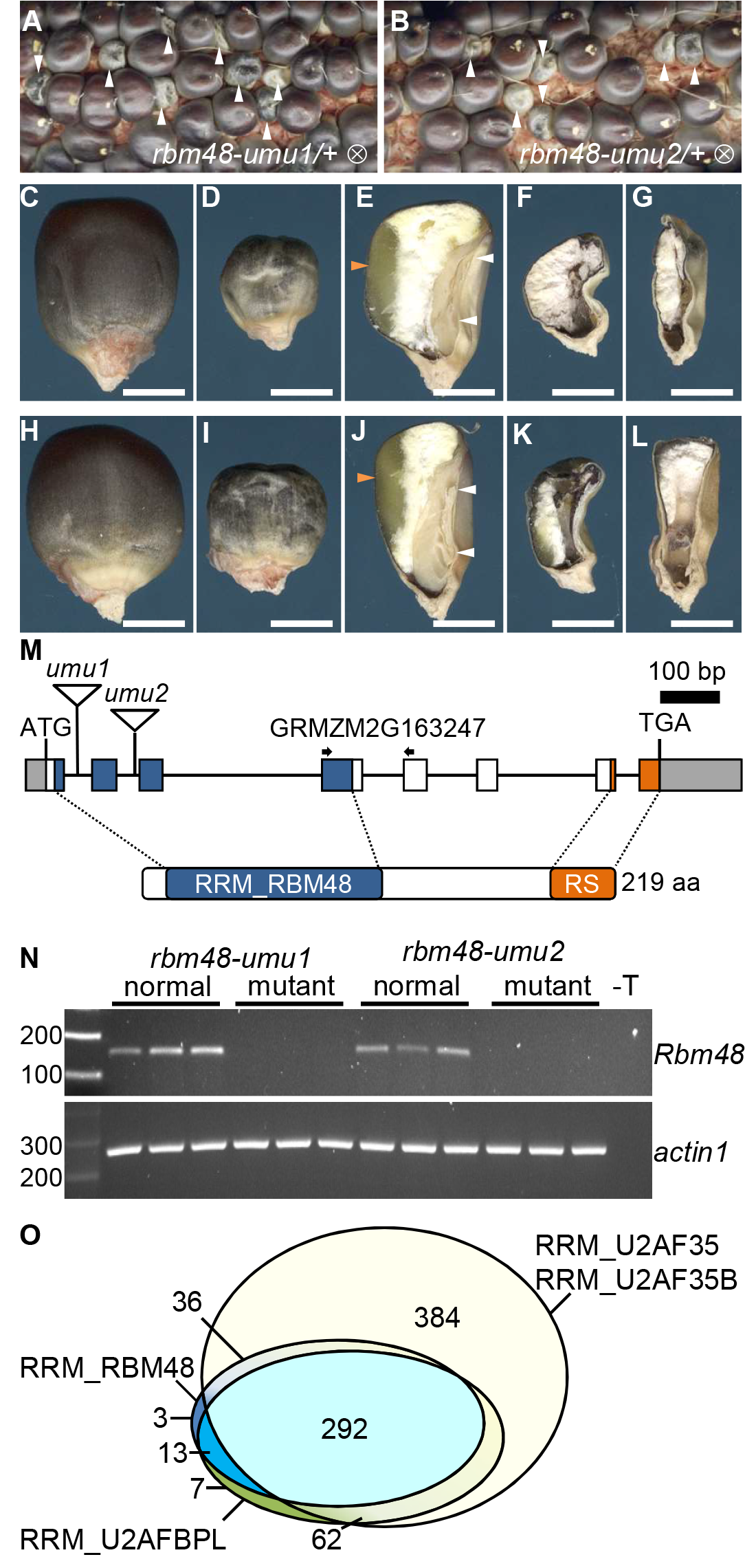
Mutant alleles of *rbm48*. (A-B) Segregating self-pollinated ears for *rbm48-umu1* and *rbm48-umu2* alleles. Arrowheads indicate *rbm48* mutant kernels. (C-L) Mature kernel phenotypes for *rbm48-umu1* (C-G) and *rbm48-umu2* (H-L). Abgerminal view of normal siblings (C,H), *rbm48-umu1* (D), and *rbm48-umu2* (I). Sagittal sections of normal siblings (E,J), *rbm48-umu1* (F, G), and *rbm48-umu2* (K, L). Scale bars are 0.5 cm in all panels. White arrowheads indicate shoot and root of the embryo. Red arrowhead indicate vitreous endosperm. (M) Schematic of the *Rbm48* locus, GRMZM2G163247, and protein domain structure. Triangles indicate transposon insertions causing *rbm48-umu1* and *rbm48-umu2*. Arrows indicate primers for RT-PCR in panel N. (N) RT-PCR of *Rbm48* and *actin1* control in *rbm48-umu1* and *rbm48-umu2*, and their normal siblings. T is a no template DNA negative control. (O) Proportional Venn diagram showing the number of species in the NCBI Conserved Domain Database with RRM domains from RGH3 (RRM_U2AFBPL), RBM48 (RRM_RBM48), and U2AF1 (RRM_U2AF35 union with RRM_U2AF35B).

Mature *rbm48* kernels have reduced endosperm size and embryos typically fail to develop (Figure 1A-L). Consistent with these morphological defects, mutant kernel composition is affected with reduced oil, starch, and seed density (Supplementary Figure 2). A small fraction of *rbm48* kernels have a less severe phenotype and develop a viable embryo that germinates. Mutant seedlings developed only 1-2 narrow leaves, stunted roots, and died around 20 days after sowing (Supplementary Figure 2). Thus, both *rbm48* alleles are lethal mutations.

RT-PCR of normal *Rbm48* cDNA from etiolated roots and shoots of W22, B73, and Mo17 inbred seedlings identified a common transcript isoform coding for a 219 aa protein (Figure 1M, Supplementary Figure 3). Alternatively spliced isoforms had premature termination codons that are predicted to be targets of nonsense mediated decay (Shaul, 2015). The predicted RBM48 protein has an N-terminal RNA recognition motif (RRM) specific to the RBM48 protein family and a 30 amino acid C-terminal RS-rich motif (Figure 1M, Supplementary Figure 3). This domain structure is common for SR proteins involved in pre-mRNA splicing (Graveley, 2000). We were not able to detect *rbm48* transcripts in mutant seedlings for either allele, and we infer that the two alleles are likely null mutations (Figure 1N).

Based on the NCBI Conserved Domains Database (Marchler-Bauer et al., 2017), the RRM domain of RBM48 is found in 344 eukaryotic species (Figure 1O). The RRM domains of RBM48 and ZRSR2/RGH3 appear to be coselected with 89% of species having an RBM48 domain also containing a RGH3/ZRSR2 RRM domain. By contrast, 50% of species with an RRM domain from the core U2 splicing factor, U2AF1, lack both ZRSR2 and RBM48 RRM domains. Like RGH3, the RBM48 RRM domain is not found in model organisms that have lost MIGs and the U12 splicing machinery. Phylogenetic analysis of 16 representative species shows the distribution of RBM48 orthologs across multiple eukaryotic kingdoms with RBM48 absent in clades lacking a U12 spliceosome including algae, nematodes, and slime molds (Figure 2). In *Arabidopsis thaliana* and *Arabidopsis lyrata*, a second copy of the RBM48 domain is fused to a predicted chloroplast-localized pentatricopeptide repeat protein (Supplementary Figure 3). This gene fusion is limited to the Arabidopsis genus. Interestingly, *D. melanogaster* has a divergent RBM48-like gene (CG34231) and a highly reduced number of MIGs. These phylogenetic data suggest a potential role for RBM48 in U12 splicing.

**Figure 2.**
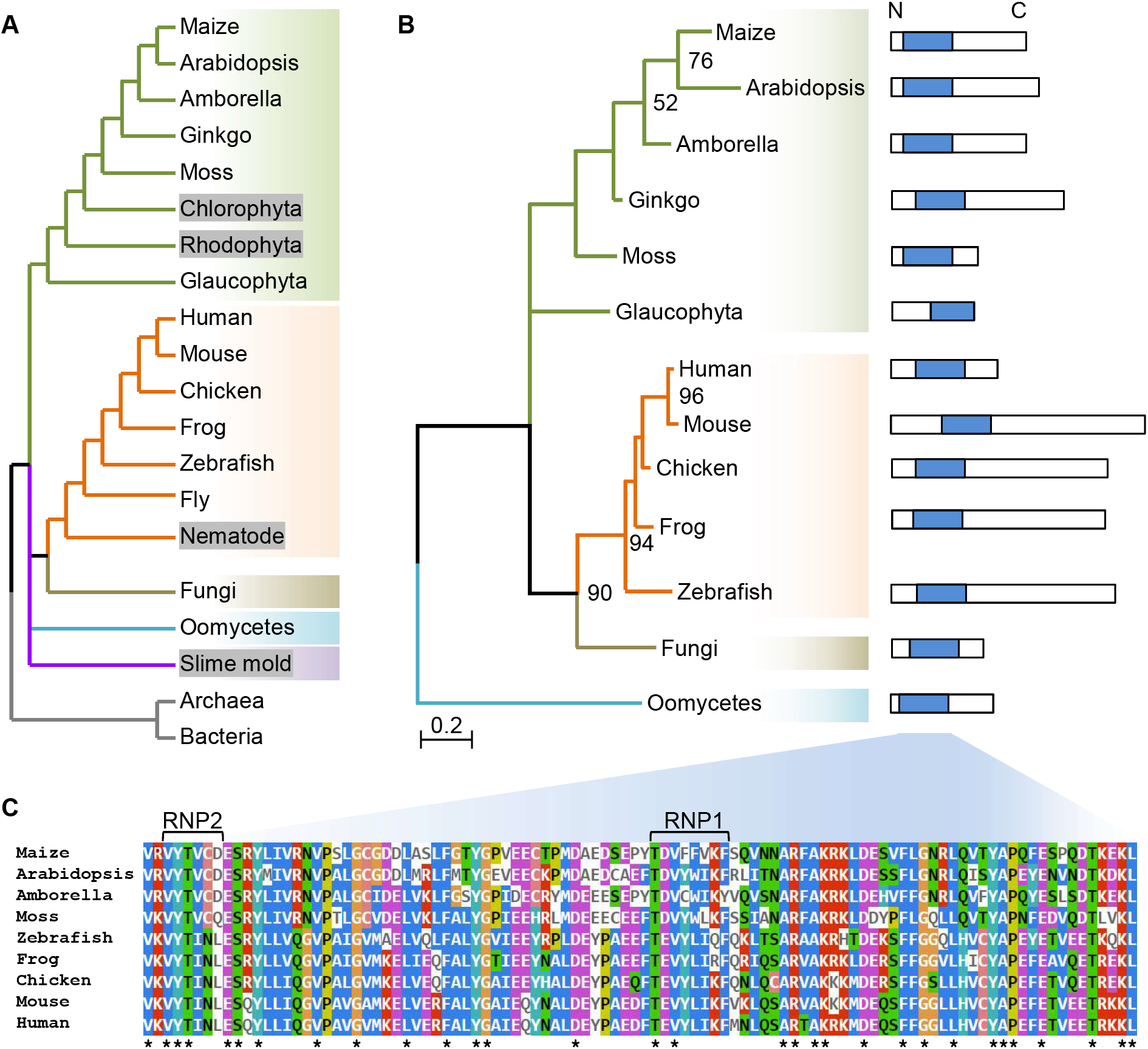
RBM48 is missing in some eukaryotic clades. (A) Species tree including significant eukaryotic model organisms. Grey boxes indicate lineages that have lost U12-type introns. *Drosophila melanogaster* (fly) encodes a hypothetical gene with an atypical RBM48 RRM-like domain. (B) Maximum likelihood tree of the RBM48_RRM domain. Bootstrap values ≥50 are reported in the corresponding nodes. Protein schematics show the RBM48 domain in blue. The glaucophyta hypothetical RBM48 protein is from *Cyanophora paradoxa*; the fungi hypothetical RBM48 protein is from *Rhizophagus irregularis* DAOM 181602; the oomycetes hypothetical RBM48 protein is from *Phytophthora sojae*. (C) Multiple sequence alignment of representative RBM48 domains. Asterisks indicate completely conserved sites. Ribonucleoprotein domains (RNP1, RNP2) of the predicted β1- and β3- seets in the RRM are indicated.

### Aberrant splicing of U12-type introns in *rbm48*

RNA splicing was assessed with mRNA-seq comparing mutant and normal sibling endosperm RNA from both *rbm48* alleles. To quantify the extent of intron splicing defects, we calculated the percent spliced out (PSO) of individual introns using exon-exon junction and intron reads (Supplementary Table 3). PSO compares the density of exon-exon junction reads and intron reads for each intron (Figure 3A). A PSO of 100 indicates complete splicing of an intron, and a value of 0 indicates all expressed transcripts retain the intron. The difference in PSO values for individual introns between normal siblings and *rbm48* mutants (ΔPSO) identifies introns that are differentially impacted. The major U2-type introns of the B73_v2 filtered gene set are largely unaffected with only 3-5% of introns in expressed genes have a ∆PSO >20% (Figure 3B-E). By contrast, 65% and 53% of U12-type introns in *rbm48-umu1* and *rbm48-umu2*, respectively, have a ∆PSO >20% suggesting more than half of U12-type introns are retained in *rbm48* mutants. Hierarchical clustering of U12-type introns based on PSO values for normal and *rbm48* mutant alleles show highly overlapping effects on intron retention, except that the *rbm48-umu1* RNA-seq experiment had a slightly stronger magnitude of defects (Figure 3F). Environmental variance between segregating ears or normal sibling contamination of the *rbm48-umu2* mutant samples may account for differences in the magnitude of defects observed.

**Figure 3.**
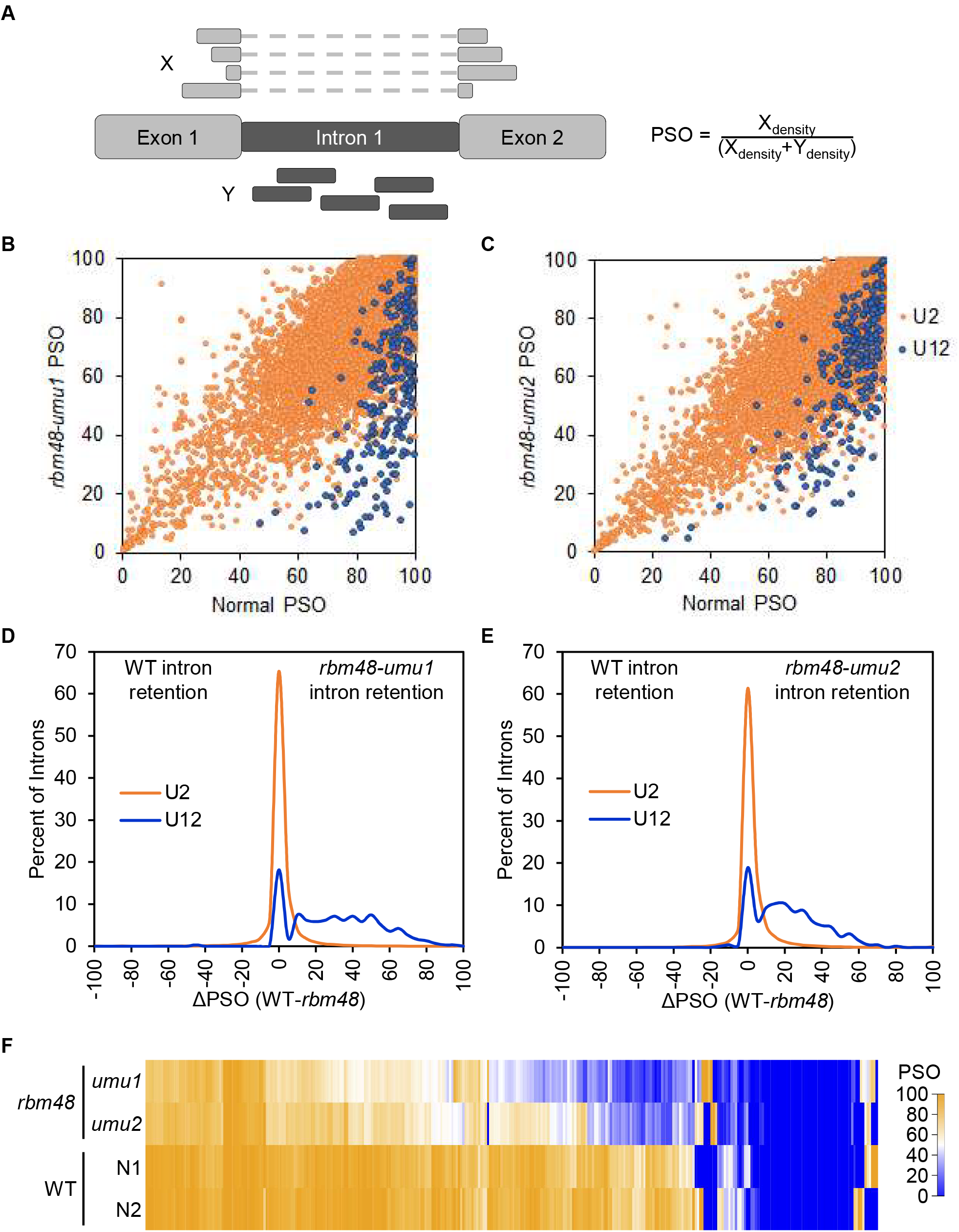
Distribution of intron retention in *rbm48* mutants. (A) Schematic of reads used to calculate PSO. (B-C) Scatter plots of mutant and normal sibling PSO values for all introns with at least 10 exon-exon junction reads in *rbm48-umu1* (B) and *rbm48-umu2* (C) mRNA-seq experiments. (D-E) Distribution of RNA splicing differences for U2-type and U12-type introns in *rbm48-umu1* (D) and *rbm48-umu2* (E) 16-18 DAP endosperm. Plots show the distribution of ΔPSO values for all introns with sufficient reads to calculate ∆PSO. Positive values have intron retention in mutant endosperm. (F) Heat map of PSO values of all U12-type introns for both *rbm48* mutant alleles and normal (N) sibling controls.

For example, the U12-type intron in GRMZM2G131321 has a ΔPSO of 65% and 50% for *rbm48-umu1* and *rbm48-umu2*, respectively. There is a high density of U12-type intron reads in the mutant RNA-seq data (Figure 4A). The U12-type intron in GRMZM2G097568 had lower ∆PSO values with a minimum of 23% for *rbm48-umu2*. No exon-exon junction reads were detected for the U12-type intron of GRMZM2G052178 in either mutant or normal samples, yet a higher density of intron reads are readily apparent in *rbm48* mutants for both genes. Semi-quantitative RT-PCR assays spanning these U12-type introns shows splicing defects with increased intron retention in both endosperm and shoot tissues (Figure 4B). In all cases, both *rbm48* alleles appear to have similar relative ratios of splice variants amplified indicating equivalent defects. RT-PCR assays confirmed eleven U12-type intron splicing defects (Figures 4B, 5E, Supplementary Figures 4-5).

**Figure 4.**
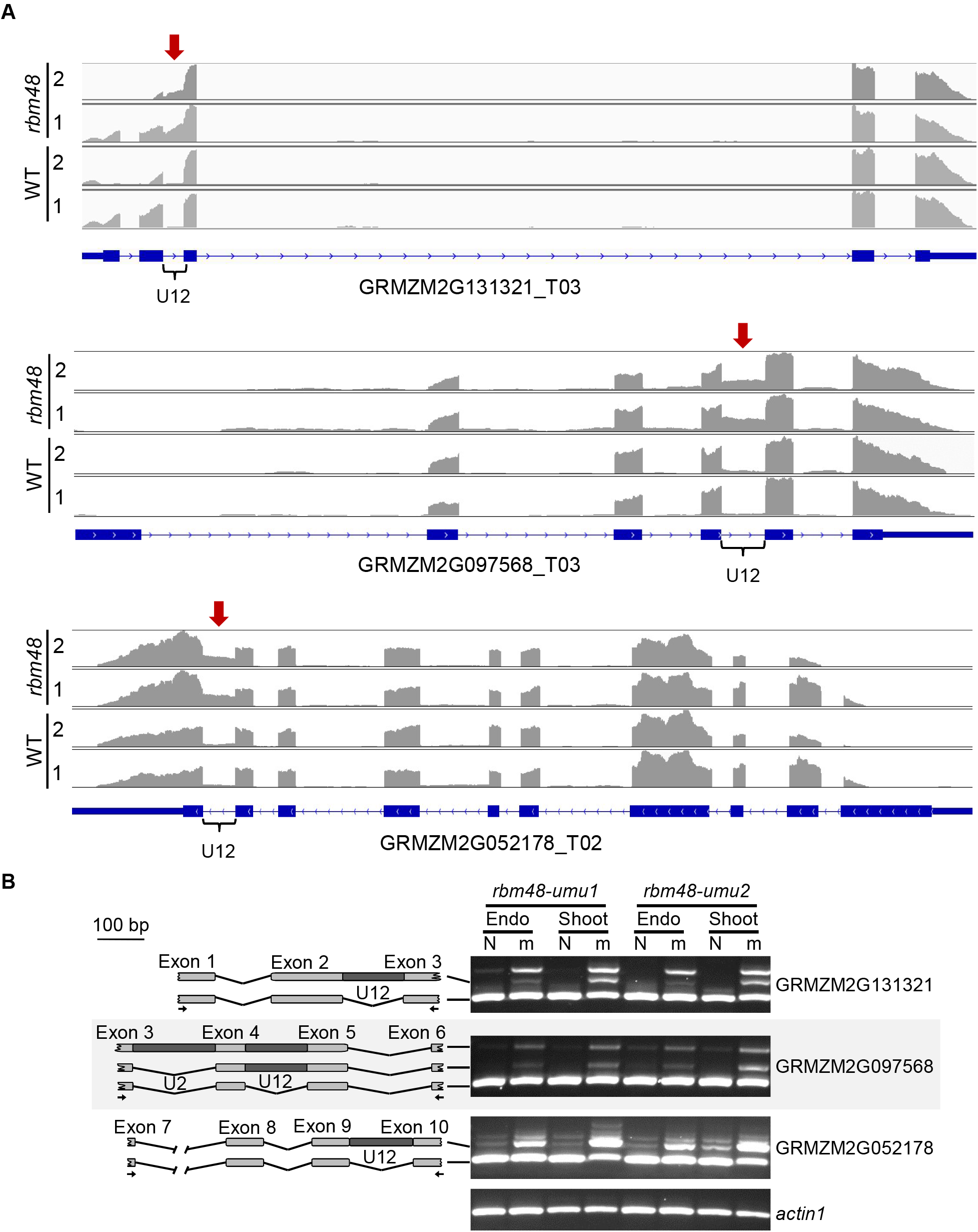
U12-type intron retention in *rbm48* mutants. (A) RNA-seq read depth of GRMZM2G131321, GRMZM2G097568, and GRMZM2G052178, homologs of human SPCS2, TRAPPC2, and BYSL, respectively. Each panel is the summed read depth of four biological replicate libraries of mutant and normal sibling (WT) endosperm tissues for *rbm48-umu1* and *rbm48-umu2* segregating ears. Red arrow and brace symbol indicate the U12-type intron with increased read depth in *rbm48* mutants. (B) RT-PCR of normal (N) and *rbm48* mutant (m) RNA from endosperm and seedling shoot tissues. Schematics show amplified products with PCR primers indicated by arrows.

**Figure 5.**
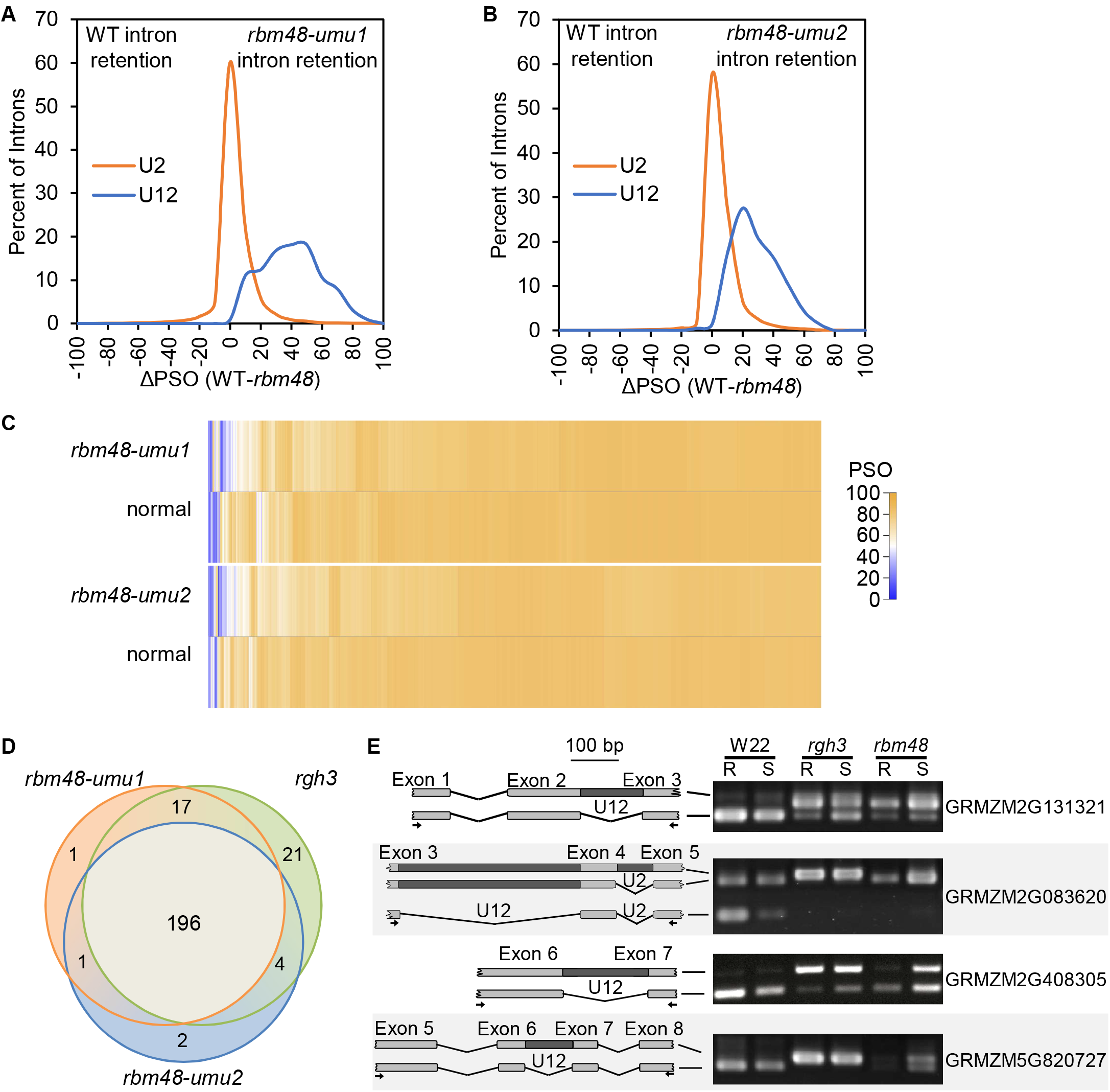
Statistically-defined splicing defects in *rbm48* mutants overlap with *rgh3*. (A-B) Distribution of ∆PSO values for introns with significant (<0.05 FDR) Fisher’s Exact Test statistics for *rbm48-umu1* (A) and *rbm48-umu2* (B). (C) Heat maps showing PSO values for U2-type introns with significant Fisher’s Exact Test statistics. (D) Venn diagram of U12-type introns that are significantly retained in *rbm48-umu1*, *rbm48-umu2*, and *rgh3* RNA-seq experiments. Introns that were tested in all three experiments were compared. (E) RT-PCR of root (R) and shoot (S) tissues from W22 inbred, *rgh3*, and *rbm48-umu1* seedlings. Schematics show amplified products with PCR primers indicated by arrows. Gene names are from the maize B73_v2 annotation.

Fisher’s exact tests of intron and exon-exon junction read counts were used to identify statistically significant splicing differences for individual introns (Supplementary Table 3). Approximately 63% of U12-type introns tested had a significant retention in both alleles (FDR<0.05). U2-type introns were largely unaffected with only 6.7% of U2-type introns having significant differences in both alleles. The median ΔPSO in these significant U2-type introns is 3%, while mis-spliced U12-type introns show a median ΔPSO of 33% (Figure 5A-C).

To identify U2-type introns with large magnitude splicing defects, we filtered the 7,535 U2-type and 201 U12-type significant introns for consistent splicing defects by requiring the ∆PSO values to be within 2-fold for both alleles and requiring at least one allele to have a |∆PSO| ≥25%. More than 68% of significant U12-type introns (43% of all U12-type introns tested) and only 4.6% of significant U2-type introns (0.26% of all U2-type introns tested) met these criteria. These larger magnitude effects included 73 U2-type introns that were spliced with higher efficiency in *rbm48* compared to normal and 279 predicted U2-type intron retention events. Twenty of the U2-type intron retention defects were in MIGs. These data indicate *rbm48* mutants confer large magnitude effects on >40% of U12-type introns with rare (~0.2%) U2-type splicing defects.

We also compared the two *rbm48* alleles with prior mRNA-seq analysis of *rgh3* (Gault et al., 2017). Of the 242 U12-type introns tested in all three mutant alleles, 196 (77%) were significantly affected in all experiments (Figure 5D). Only 24 (10%) of the tested U12-type introns were significant in only one of the *rbm48* alleles, which is within the expected false positive limits of the FDR correction for multiple testing. Consequently, there is no statistical difference between the U12-type introns affected in either *rbm48* allele. Direct RT-PCR comparisons of mutant seedling tissues confirmed similar U12 splicing defects in *rgh3* and *rbm48-umu1* (Figure 5E, Supplementary Figure 5). These analyses indicate that *rgh3* and *rbm48* have extensive overlap in U12 splicing defects.

### *rbm48* disrupts endosperm cellular development

Differential gene expression analysis of the *rbm48* RNA-seq data identified large scale changes. Of 16,745 transcripts expressed at least one transcript per million (TPM) in one genotype, 5,771 transcripts were differentially expressed (FDR <0.05, >2-fold change) in either *rbm48-umu1* or *rbm48-umu2* (Figure 6A-B, Supplementary Table 4). There were 1,811 up-regulated and 766 down-regulated transcripts in both alleles. A subset of these gene expression differences were confirmed with semi-quantitative RT-PCR (Figure 6C). GO term enrichment analysis identified nutrient reservoir activity as the most significantly enriched term among differentially expressed genes (Supplementary Table 5).

**Figure 6.**
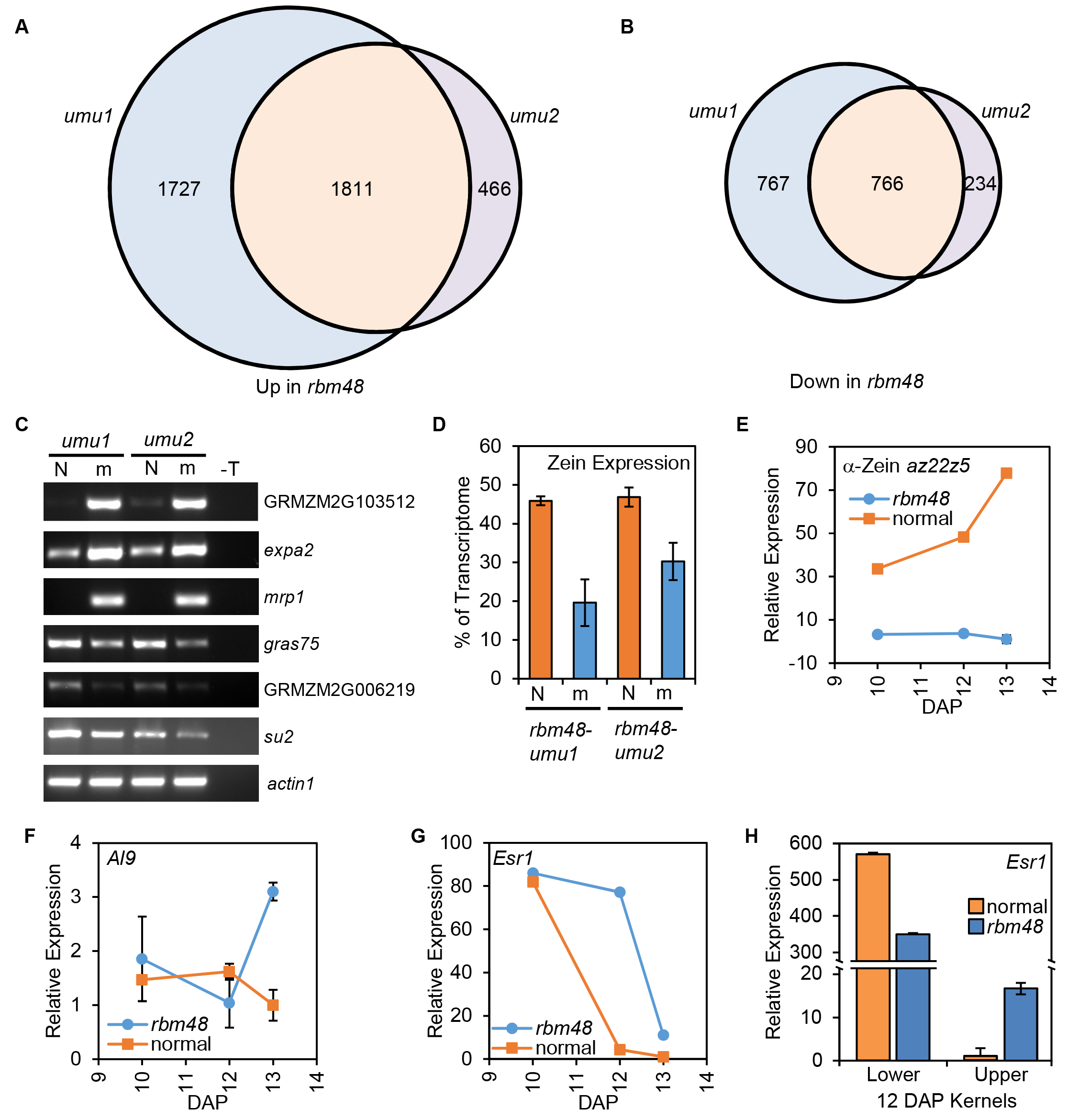
Expression defects in *rbm48*. (A-B) Venn diagrams showing the overlap of up- (A) ans down-regulated (B) DEGs in *rbm48-umu1* and *rbm48-umu2.* (C) RT-PCR of normal (N) and *rbm48* mutant (m) endosperm RNA used for RNA-seq. GRMZM2G103512, *expa2*, and *mrp1* were predicted to have increased levels in *rbm48* mutants, while *gras75*, GRMZM2G006219, and *su2* were predicted to show decreased expression in *rbm48*. The *actin1* locus was used as a loading control. T is a no cDNA template negative control. Gene names are MaizeGDB locus identifiers. (D) Proportion of total transcriptome represented by 39 zein transcripts based on endosperm RNA-seq. Error bars are standard deviation of four biological replicates. (E-G) Quantitative RT-PCR of total RNA extracted from whole kernels in *rbm48* and normal siblings for *α-Zein* (E), *Al9* (F), and *Esr1* (G). Error bars are standard deviation of three biological replicates. (H) Quantitative RT-PCR of *Esr1* expression in total RNA extracted from lower and upper half kernels. Error bars are standard deviation of three biological replicates.

Nearly 80% the nutrient reservoir activity genes affected are seed storage proteins, such as zeins. At this stage of endosperm development, zein expression accounts for 45% of the total transcriptome in the normal sibling endosperm and only 20-30% of the total transcriptome in *rbm48* mutants (Figure 6D). We confirmed that α-zein expression was reduced in *rbm48-umu1* using qRT-PCR in a developmental series of whole kernel RNA (Figure 6E). The *rgh3* mutant also reduces and delays zein accumulation in the endosperm as part of an overall disruption of endosperm cell differentiation (Fouquet et al., 2011).

A focused analysis of endosperm cell type marker genes revealed that cell-type specific genes had aberrant expression from 10-13 days after pollination (DAP). Cell-type markers for the aleurone (*Al9*), embryo surrounding region (*Esr1*), and basal endosperm transfer cell layer (*Betl2*, *Tcrr1*) had increased expression late in development (Figure 6F-G, Supplementary Figure 6).

Many of these whole kernel expression patterns were strikingly different from those observed from the lower half of kernels analyzed in the *rgh3* mutant (Fouquet et al., 2011). For example, *Esr1* is expressed at a basal level in the lower half of *rgh3* mutants, while *Esr1* is up regulated in whole *rbm48* kernels. Extractions of the upper and lower half of 12 DAP kernels showed that this discrepancy is due to reduced expression of *Esr1* in the lower half of the kernel, where *Esr1* is normally expressed, and increased expression of *Esr1* in the upper half of the kernel (Figure 6H). Quantitative expression analysis of the lower half of 12 DAP kernels also showed that, while *Al9* is induced, *Betl2* and *Esr1* are reduced consistent with similar observations of *rgh3* (Supplementary Figure 6). These results indicate that *rbm48* endosperm cells fail to express cell type specific markers in the correct domains of the kernel.

Cellular phenotypes of *rbm48* endosperm are also consistent with cell differentiation defects. Transverse sections of 12 DAP kernels show that *rbm48* starchy endosperm cells are smaller than normal siblings (Figure 7A-B). At this stage, the basal endosperm transfer layer (BETL) is differentiated in normal kernels, visible as elongated cells with intense Schiff’s staining due to secondary cell wall ingrowths (Figure 7C). The BETL in *rbm48* is not developed into elongated cells (Figure 7D). Cell-type specific reporter lines indicate that the *rbm48* BETL region differentiates into aleurone. A GUS reporter gene driven by the *Viviparous1* promoter (pVP1:GUS) marks ABA responsive cells including the aleurone and embryo, while the *Betl1* promoter (pBetl1:GUS) marks BETL cells. These reporter transgenes have aberrant expression in *rbm48-umu1* mutants with pVp1:GUS expressing in the basal epidermal endosperm and pBetl1:GUS having low expression in the same region (Figure 7E-H, Supplementary Figure 7). These data support the model that epidermal endosperm cells express aleurone markers regardless of position in the endosperm.

**Figure 7.**
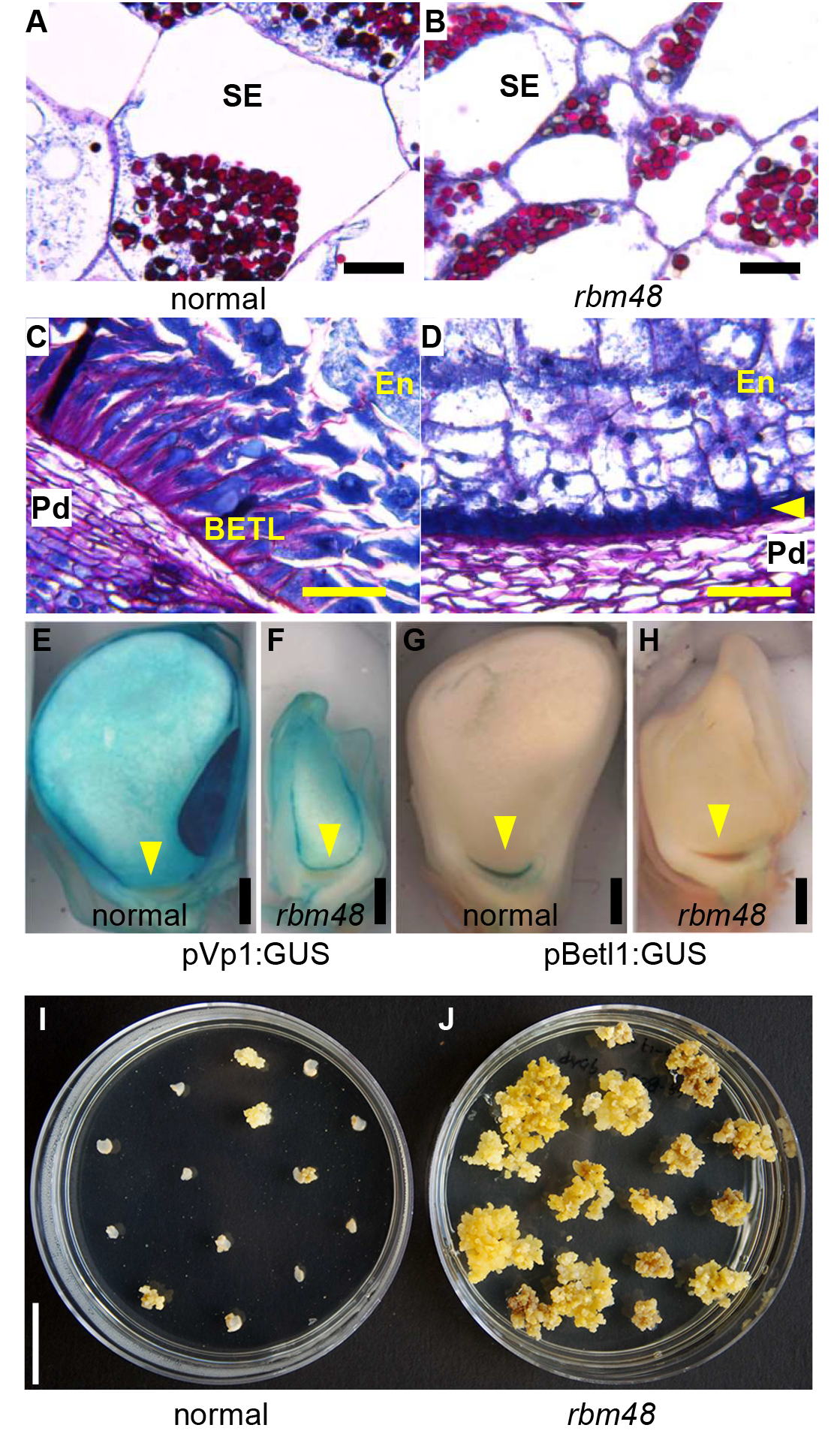
Endosperm cell differentiation defects in *rbm48*. (A-D) Transverse sections of 12 DAP normal sibling (A,C) and *rbm48-umu1* (B,D) kernels stained with Schiff’s reagent and aniline blue-black. Insoluble carbohydrates in cell walls and starch grains stain fuchsia; nucleoli, nuclei, and cytoplasm stain different intensities of blue. Scale bar is 50 μm. (A-B) Internal starchy endosperm cells (SE). (C-D) BETL region with inner endosperm (En) and maternal pedicel (Pd). Yellow arrowhead indicates small epidermal cells in the BETL region. Scale bar is 50 μm. (E-H) Maize transgene reporter lines for *Vp1* (pVp1:GUS) and *Betl2* (pBetl2:GUS) in normal siblings (E,G) and *rbm48-umu1* mutants (F,H) at 13 DAP. Blue stain indicates GUS expression. Yellow arrowhead indicates the BETL region. Scale bar is 1 mm. (I-J) Tissue culture response of 9 DAP endosperm from normal (I) sibling and *rbm48* (J) seeds. Plates show growth response after 35 days of culture. Scale bar is 2 cm.

Mutant *rgh3* endosperm retains the ability to proliferate in cell culture at late time-points in seed development (Fouquet et al., 2011). Endosperm tissue culture responses for *rbm48-umu1* mutants and normal siblings were tested. At 6 DAP, normal and mutant endosperm grow equally well in the culture assay, but at 9-18 DAP, only *rbm48* endosperms are able to grow as a callus in the culture assay (Figure 7I-J, Supplementary Figure 7). Thus, *rbm48* mutants have prolonged endosperm cell proliferation and delayed or aberrant differentiation, similar to *rgh3* mutants.

### RBM48 interacts with U12 and U2 splicing factors

The evolutionary coselection, U12 splicing, and endosperm cell differentiation defects in *rbm48* and *rgh3* mutants raises the possibility that the proteins interact. We investigated the subcellular localization of RBM48 with GFP fusion proteins that were transiently expressed in *Nicotiana benthamiana* leaves. Full length RBM48 is localized to the nucleus and is associated with the nuclear speckles (Figure 8A). Deletion of the RRM domain still localized the protein in nuclear speckles, albeit at a lower expression level, while deletion of the C-terminus of RBM48 resulted in diffuse cytoplasmic signal (Figure 8B-C). These results are consistent with the localization of SR proteins in both plants and animals to sub-nuclear speckles via the RS domain (Mori et al., 2012; Saitoh et al., 2012; Tillemans et al., 2005; Tillemans et al., 2006; Cazalla et al., 2002).

**Figure 8.**
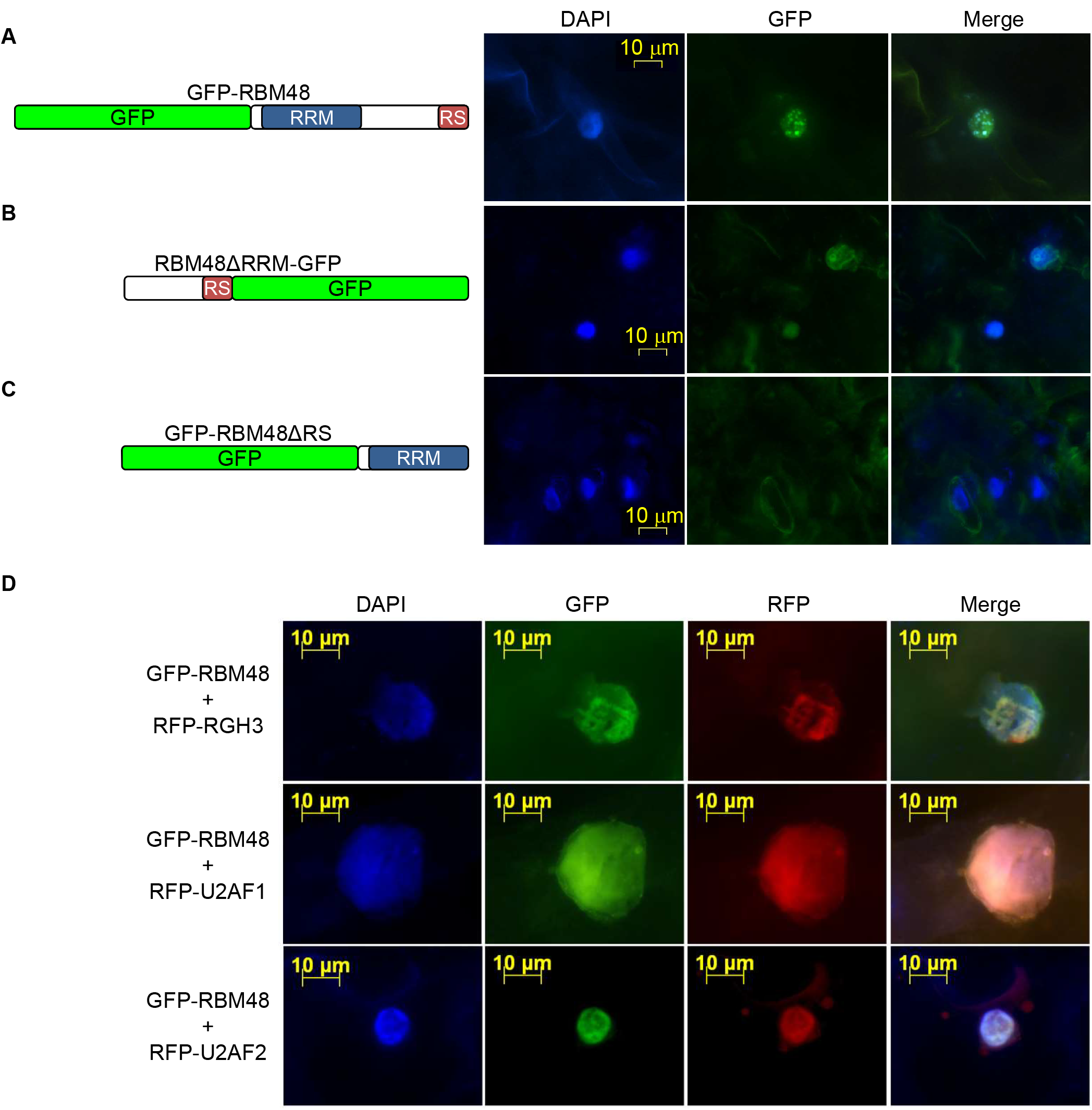
RBM48 localizes to the nucleus with U2AF and RGH3. (A-C) Subcellular localization of RBM48 and RBM48 domain deletions. (A) N-terminal GFP fusion of the full length RBM48 protein. (B) C-terminal GFP fusion deleting the RBM48_RRM domain. (C) N-terminal GFP fusion deleting the C-terminal 109 amino acids including the RS domain. (D) Transient co-expression of GFP-RBM48 with RFP-tagged RGH3, U2AF1, and U2AF2 in *N. benthamiana* leaves. DAPI staining of DNA marks nuclei in blue.

Co-expression of GFP-RBM48 with RFP-tagged RGH3 or the U2AF1 and U2AF2 subunits of U2AF resulted in extensive overlap of fluorescent signals (Figure 8D). Human U2AF2 interacts with U2AF1 and the RGH3 ortholog, ZRSR2 (Tronchere et al., 1997; Kielkopf et al., 2001). The co-localization of RBM48 with these factors suggests that RBM48 may interact in a larger complex. Human U2AF is involved in the selection of 3’ splice sites of U2-type introns, while ZRSR2 binds the 3’ splice sites of U12-type introns (Tronchere et al., 1997; Guth et al., 2001; Shen et al., 2010).

Bimolecular fluorescence complementation (BiFC) assays of RBM48 with RGH3, U2AF1, and U2AF2 support the model of a larger protein complex of these factors. BiFC tests for fluorescence complementation of split, non-fluorescent fragments of YFP that are brought in close vicinity by interacting fusion proteins (Citovsky et al., 2006; Kerppola, 2008). The N-terminal or C-terminal fragments of YFP were fused with RBM48 and transiently co-expressed in different pairwise combinations with N- or C-terminal YFP fusion constructs of RGH3, U2AF1, and U2AF2 in *N. benthamiana* leaves. All combinations of RBM48 with full length RGH3, U2AF1, and U2AF2 resulted in YFP signal indicating these proteins are in close proximity in plant cells (Figure 9). An inframe deletion of the RGH3 RRM domain (RGH3ΔUHM), also known as the U2AF homology motif (UHM), fails to complement YFP indicating that the UHM domain is needed for RBM48 and RGH3 association (Figure 9A). When expressed with U2AF2 fusions, the RGH3∆UHM construct results in BiFC signal showing that the construct is capable of reporting close proximity of proteins in the nucleus (Gault et al., 2017).

**Figure 9.**
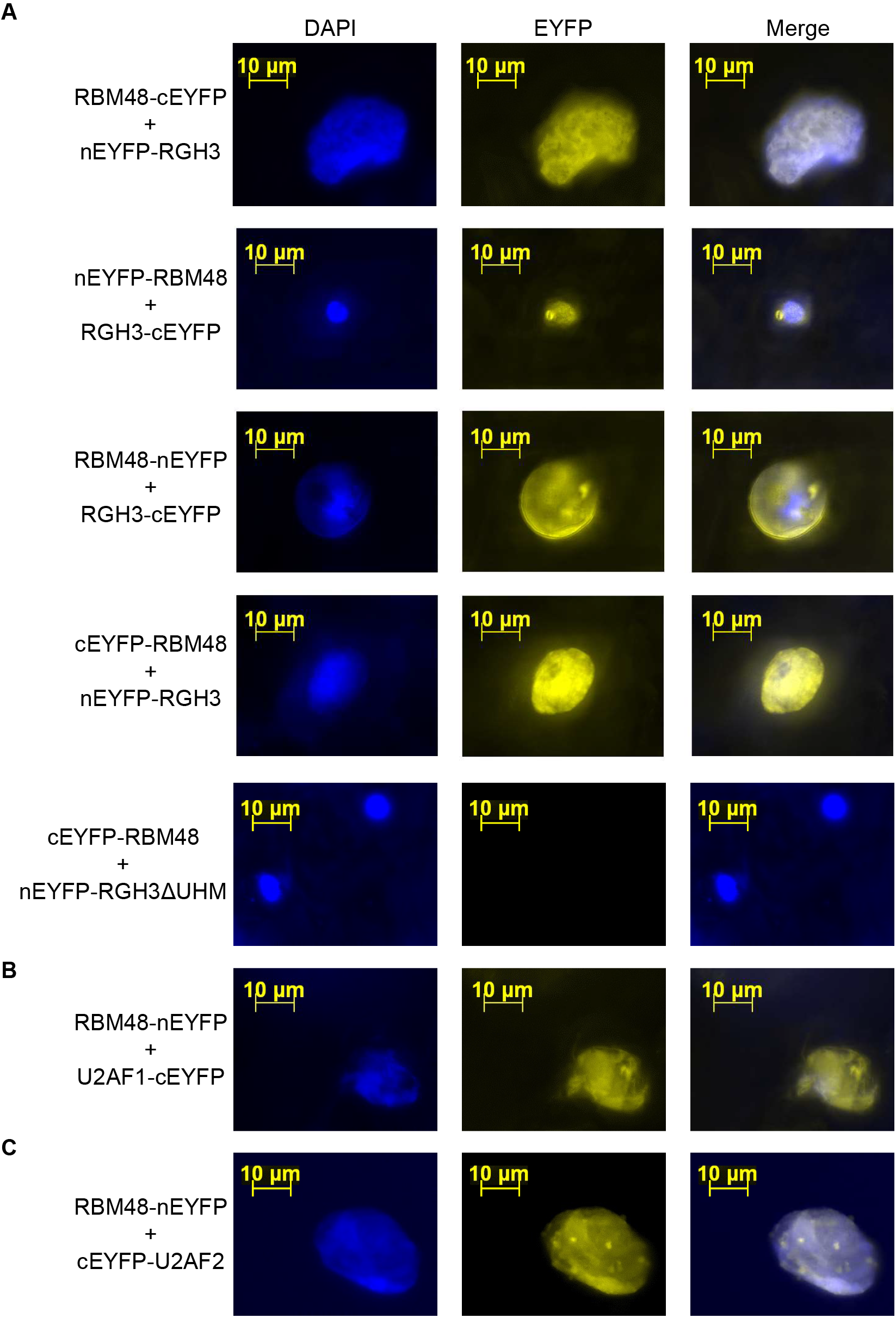
BiFC interactions of RBM48 with RGH3, U2AF1, and U2AF2. BiFC assays showing transient co-expression in *N. benthamiana* leaves of fusion proteins with N-terminal EYFP (nEYFP) or C-terminal EYFP (cEYFP). Yellow signal indicates reconstituted EYFP. DAPI staining of DNA marks nuclei in blue. (A) Co-expression of four combinations of RBM48 and RGH3 N- and C-terminal fusion proteins as well as the RGH3ΔUHM in-frame domain deletion. (B) Co-expression of RBM48 and U2AF1 fusions. (C) Co-expression of RBM48 and U2AF2 fusions.

We then tested direct protein-protein interactions with *in vitro* pull-down assays. Recombinant GST-His-RBM48 fusion protein was immobilized to glutathione magnetic beads and incubated with *E. coli* lysate containing recombinant His-tagged RGH3, U2AF1, U2AF2, or Armadillo Repeat Containing7 (ARMC7, GRMZM2G106137). Human RBM48 interacts with ARMC7 (Hart et al., 2015). In all cases, RBM48 can purify the His-tagged proteins from bacterial lysates (Figure 10A). Moreover, GST-His-RGH3 pulls down His-tagged U2AF1 and U2AF2, but not Cbes_2310, a protein originating from *Caldicellulosiruptor bescii* (Figure 10B). These results support conserved interactions in maize for RBM48 with ARMC7 and RGH3 with U2AF2. In addition, we found that RBM48 and RGH3 interact with each other as well as with U2AF1.

**Figure 10.**
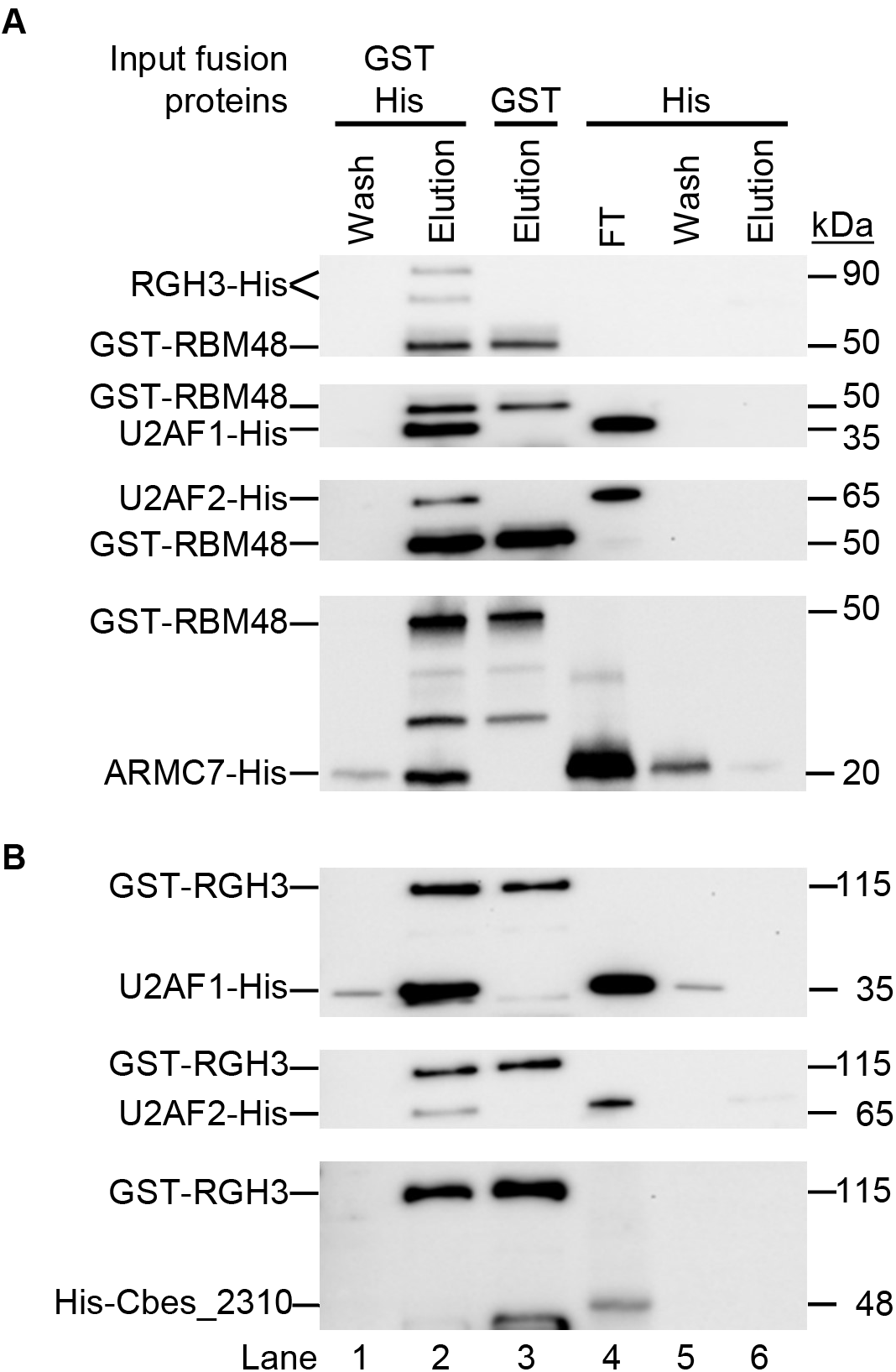
Direct protein-protein interactions with RBM48. (A) Western blots of *in vitro* pull down assays with GST-His-tagged RBM48. GST-His-RBM48 immobilized on glutathione beads was incubated with bacterial lysate containing His-tagged RGH3, U2AF1, U2AF2, or ARMC7. (B) Western blots of *in vitro* pull down assays with GST-His-RGH3 immobilized on glutathione beads and bacterial lysates containing His-tagged U2AF1, U2AF2, or Cbes_2310. Recombinant proteins are detected with αHis monoclonal antibodies. Lanes 1-2 show the wash and elution of the pull down assay with both recombinant proteins. Lane 3 is the elution of the GST-His-tagged protein alone. Lanes 4-6 show the level of binding of the His-tagged protein with unbound glutathione beads. FT is the flow through lysate after incubation with unbound beads.

## Discussion

### RBM48 is a U12 splicing factor

Genetic and molecular analysis of maize *rbm48* revealed extensive defects in U12-type intron splicing with more than 60% of MIGs significantly mis-spliced. Although RNA-seq analysis showed a greater magnitude of U12-type intron splicing defects in *rbm48-umu1* (Figures 3 and 5), we did not find statistically significant different sets of U12-type introns affected in the two alleles of *rbm48*. Moreover, kernel and seedling phenotypes of the two alleles are very similar at qualitative and quantitative levels (Figure 1, Supplementary Figure 2). Both alleles show a range of kernel phenotypes throughout development, and it is very likely that environmental differences explain much of the difference in magnitude of U12 splicing defects observed for the two alleles in the RNA-seq experiment.

Defects in U2 splicing were also observed in a fraction of introns, but these are predominantly of small magnitude with >95% of significantly affected U2-type introns splicing at ±25% of normal levels. Defects in MIG splicing have very similar patterns in both *rbm48* and *rgh3* with adjacent U2-type introns being mis-spliced in a subset of MIGs. Mis-splicing of neighboring U2-type introns in minor splicing mutants has been observed in multiple species (Madan et al., 2015; Horiuchi et al., 2018). These data indicate that maize RBM48 is a minor splicing factor. No RNA processing role has been assigned yet to RBM48, and the human ortholog was not identified in biochemical purification of U12 spliceosomes (Schneider et al., 2002; Will et al., 2004). However, RBM48 is conserved among organisms that have a U12 spliceosome and has been coselected with ZRSR2/RGH3 throughout eukaryotic evolution. Based on this evolutionary conservation and our evidence that maize RBM48 is a U12 splicing factor, we predict that animal RBM48 proteins will also function in U12 splicing.

### RBM48 interacts with RGH3, U2AF, and ARMC7

The protein-protein interactions we found between RBM48 with RGH3, U2AF, and ARMC7 may help explain the impact on U2-type introns in a subset of U12 splicing mutants (Verma et al., 2017). A low frequency of U2-type intron splicing defects have been observed for mutations in ZRSR2 homologs and RNPC3 (Madan et al., 2015; Gault et al., 2017; Horiuchi et al., 2018; Argente et al., 2014). RGH3/ZRSR2 and RNPC3 are required for U12-type intron recognition (Frilander and Steitz, 1999; Shen et al., 2010). Human ZRSR2 was known to interact with U2AF2 (Tronchere et al., 1997), but we found more extensive connection between U12 splicing factors and U2AF. Both U2AF and ZRSR2 are responsible for 3’ splice site recognition of U2-type and U12-type introns, respectively (Shen et al., 2010). By contrast, U2-type splicing defects have not been reported in human mutant alleles of U4atac, which are expected to recognize U12-type introns but not complete splicing reactions (Verma et al., 2017). The interaction data and the presence of a low frequency of U2-type splicing defects suggests that RBM48 may also participate in U12-type intron recognition. A direct test of this hypothesis requires an *in vitro* RNA splicing assay, which has only recently been developed for the major spliceosome in plants (Albaqami and Reddy, 2018).

Our study also illustrates the importance of genetic analysis of RNA splicing factors to determine RNA processing functions. Recently, mutant models for several splicing factors including ZRSR2, ZRSR1, SMN, and FUS showed a predominant *in vivo* function in minor intron splicing even though the proteins are thought to have broader roles based on *in vitro* splicing assays or protein-protein interactions (Madan et al., 2015; Gault et al., 2017; Horiuchi et al., 2018; Doktor et al., 2017; Jangi et al., 2017). RNA-seq analysis of mutant transcriptomes was able to resolve the predominant defects as impacting U12-type introns.

### U12 splicing mutants in plants and animals have divergent developmental effects

In plants, genetic analysis of U12 splicing factors has defined two phenotypic syndromes. Transgenic knockdown alleles for multiple U12 splicing factors in *Arabidopsis thaliana* develop serrated leaves, delay senescence, and dwarf inflorescence branches, which can be complemented by exogenous application of gibberellins (Kim et al., 2010; Jung and Kang, 2014; Xu et al., 2016). Strong alleles of U12 splicing factors in either Arabidopsis or maize disrupt seed development (Figure 1) (Kim et al., 2010; Fouquet et al., 2011). In maize, aberrant U12-type intron splicing disrupts both endosperm and embryo development. However, reduced U12 splicing efficiency does not cause cellular lethality, because *rgh3* and *rbm48* both show increased cell proliferation in endosperm tissue culture (Figure 7) (Fouquet et al., 2011). These mutant phenotypes argue that efficient U12 splicing is required for cell differentiation rather than cell viability. Neither *rbm48* nor *rgh3* completely block U12 splicing, and it is likely that a complete block would cause cellular lethality.

Mutations disrupting U12 splicing in human cell lines reduce proliferation and are classified as core fitness genes (Hart et al., 2015; Blomen et al., 2015). Interestingly, ARMC7, an RBM48 interacting protein, was also found to be a core fitness gene in these studies. It is somewhat surprising that homologous mutations in maize *rgh3* and *rbm48* have contrasting cell proliferation defects in endosperm tissue culture (Figure 7) (Gault et al., 2017; Fouquet et al., 2011).

Although MIGs represent a small fraction of eukaryotic genes, plants and animals have MIGs in conserved cellular and biochemical pathways such as autophagy, secreted protein glycosylation, DNA damage repair, cell cycle, and histone methyltransferases (Gault et al., 2017).

Disruption of multiple cell cycle and DNA damage repair genes would be expected to promote cell proliferation. Although *rbm48* and *rgh3* promote cell proliferation, *ZRSR2* mutants reduce proliferation. However, *ZRSR2* mutations are considered drivers towards cancer (Cazzola et al., 2013). Gault et al. (2017) suggested that divergence in MIGs or divergence in the location of U12-type introns within conserved MIGs are likely causes for the arrest of animal cell proliferation versus increased plant cell proliferation. For example, a subset of E2F cell-cycle transcription factors are MIGs in both maize and humans, but the U12-type introns are in different positions relative to the open reading frames. Retention of the human U12-type intron is predicted to knockout function through nonsense mediated decay, while retention of the U12-type intron in maize does not impact RNA stability and is unlikely to disrupt protein function (Gault et al., 2017).

### Divergence of U12 splicing phenotypes likely result from fractionation of MIGs

U12-type introns have longer and more conserved 5’ splice site and branch point consensus sequences that are recognized through cooperative binding of the U11 and U12 snRNPs (Frilander and Steitz, 1999). Consequently, divergence in MIGs between eukaryotic species predominantly is due to U12-type intron loss or mutation to U2-type introns (Burge et al., 1998; Lin et al., 2010). A large fraction of MIGs have been maintained in a conserved genetic architecture between plant and animal species suggesting that U12 splicing efficiency could have similar effects on the cell (Gault et al., 2017).

Minor splicing mutants in animals have been shown to cause defects in cell differentiation. A subset of human myelodysplasia patients have mutations in *ZRSR2* that result in aberrant U12 splicing and reduced terminal differentiation of myeloid blood cell types (Madan et al., 2015). Mutation of the mouse *ZRSR1* paralog of *ZRSR2* leads to blood and sperm differentiation defects (Horiuchi et al., 2018). Patients with Roifman syndrome have mutations in the human U4atac snRNA along with defects in both myeloid and lymphoid blood cell differentiation (Heremans et al., 2018). These examples suggest a common requirement for efficient splicing of U12-type introns to promote the differentiation of a subset of eukaryotic cell types. However, minor splicing mutations cause a range of human diseases and diverse phenotypes in animal models (Horiuchi et al., 2018; Verma et al., 2017; Pessa et al., 2010; Markmiller et al., 2014). The similarity of endosperm defects in *rbm48* and *rgh3* suggest a mutant ideotype for the developmental pathways sensitive to U12-type intron splicing efficiency. Maize genetic screens for delayed endosperm cell differentiation and prolonged cell proliferation could discover both U12 splicing mutants and specific MIGs needed to promote cell differentiation.

## Methods

### Genetic stocks

The *rbm48-umu1* allele was isolated as a visual *rough endosperm* (*rgh*) mutant from the UniformMu transposon-tagging population (McCarty et al., 2005). The *rbm48-umu2* allele was identified from the UniformMu reverse genetics resource as *Mu* insertion mu1056623 in stock UFMu-07152 (McCarty et al., 2013; Settles et al., 2007). Both *rbm48-umu1* and *rbm48-umu2* were backcrossed five times into W22, B73, Mo17, and A636 for phenotypic analysis. An *rbm48-umu1* B73 x W22 F_2_ mapping population was generated by self-pollinating the F_1_ hybrid for the B73 introgression. The pBetl1:β-glucuronidase (GUS) and pVp1:GUS reporter transgenics have been described (Hueros et al., 1999; Cao et al., 2007). The *rgh3* mutant stock was maintained as a *Pr1 rgh3/pr1 Rgh3* stock in the W22 inbred background (Fouquet et al., 2011). All plant materials were grown at the University of Florida Plant Science Research and Education Unit in Citra, FL or greenhouses located at the Horticultural Sciences Department in Gainesville, FL.

### Molecular cloning of *rbm48*

The *rbm48-umu1* allele was mapped with bulked segregant analysis (BSA) using a pool of 93 homozygous *rgh* kernels from the B73 x W22 F_2_ mapping population. DNA was extracted from the pooled kernels and genotyped with 144 single nucleotide polymorphism (SNP) markers using the Sequenom MassARRAY platform at the Iowa State University Genomic Technologies Facility as described (Liu et al., 2010). Relative enrichment for the W22 genotype was calculated from non-segregating pooled samples.

*Mu* flanking sequence tags (MuFSTs) were generated from a 12 x 12 grid of 144 UniformMu *rgh* mutants including *rbm48-umu1*. Transposon-flanking sequences were amplified from pooled DNA using MuTAIL-PCR (Settles et al., 2004). The PCR products were sequenced with a Roche Genome Sequencer FLX System using custom A primers to add pool-specific barcodes and sequence from the *Mu* terminal inverted repeat (TIR). Resulting sequences were filtered for intact TIR sequences and mapped to the B73_v2 genome with blastn. Identical insertion sites sequenced from a single row and a single column pool were assigned to individual *rgh* mutant isolates. The ten unique MuFSTs in *rbm48-umu1* were co-aligned to the BSA map position to identify *rbm48-umu1* as the likely causative mutation. Co-segregation of *rbm48-umu1* with the *rgh* kernel phenotype was tested by extracting DNA from segregating kernels and amplifying the wild-type or *Mu*-insertion alleles using gene specific primers and the TIR5 primer as described (Settles et al., 2007).

### Kernel phenotypes

Segregating ears and mature kernels were imaged on a flatbed scanner. Both *rbm48* alleles were analyzed for kernel composition phenotypes by a single-kernel grain analyzer with a microbalance and near infrared spectrometer (Spielbauer et al., 2009; Gustin et al., 2013). At least 30 normal and 30 *rgh* kernels were sampled from three segregating ears of each allele. Four technical replicate kernel weight and NIR spectra were collected from each kernel. Average single-kernel composition predictions for each biological replicate were used as the kernel composition phenotypes shown in Supplementary Figure 2.

### RT-PCR

Tissues were dissected, frozen in liquid nitrogen, and ground to a powder. RNA was extracted from 100 mg of tissue and mixed with 200 μL of RNA extraction buffer (50 mM Tris-HCl, pH 8, 150 mM LiCl, 5 mM EDTA, 1% SDS in DEPC treated water). The slurry was extracted using phenol:chloroform and Trizol (Invitrogen). RNA was precipitated from the aqueous fraction using isopropanol and washed with 70% ethanol. RNA pellets were re-suspended in nuclease free water (Sigma) treated with Purelink^Tm^ DNase (Invitrogen). RNA was further purified using an RNeasy MinElute Cleanup Kit (Qiagen). M-MLV reverse transcriptase (Promega) used 1 μg total RNA as template to synthesize cDNA.

Semi-quantitative RT-PCR was carried out by modifying the methods from Bai and DeMason (2006). The PCR conditions were 3 min at 94°C, followed by 24-28 cycles of 30 sec at 94°C, 30 sec□at the appropriate annealing temperature for the primers and 1□min at 72°C, followed by a final extension of 5□min at 72°C. Quantitative RT-PCR used a StepOnePlus real-time PCR machine (Applied Biosystems) with 1X SYBR^®^ Green PCR Master Mix (Applied Biosystems) as described (Fouquet et al., 2011; Bai et al., 2016). The normalized expression level of each gene represents the average of three replicates of three biologically distinct kernel pools relative to *actin1* using the comparative cycle threshold (ΔΔ*C*_t_) method (Livak and Schmittgen, 2001). Primer sequences are Supplementary Table 6.

### Phylogenetic analysis

Coselection was detected by parsing the species names from the “specific protein” lists for the RRM_RBM48, RRM_U2AFBPL, RRM_U2AF35, and RRM_U2AF35B domains in the NCBI Conserved Domains Database (Marchler-Bauer et al., 2017) A representative set of RBM48 proteins were aligned with MUSCLE (version 3.8.31) using default parameters (Edgar, 2004). The multiple sequence alignment of the RBM48_RRM was input to RAxML (version 8.2.3) (Stamatakis, 2014) to construct a maximum likelihood phylogeny with the LG4X amino acid replacement matrix model and 1,000 bootstrap replicates (Le et al., 2012). Eight tree searches were completed to further test for tree stability. The phylogeny was rooted by the oomycetes RBM48 protein and displayed with Dendroscope (version 3.2.10) (Huson and Scornavacca, 2012).

### RNA-Seq analysis

Endosperm tissue from 16-18 DAP normal and *rbm48* kernels in the W22 genetic background was dissected and frozen in liquid nitrogen. Total mRNA was extracted from four biological replicates of paired *rbm48* mutant and normal sibling pools. Non-strand specific TruSeq^TM^ (Illumina) cDNA libraries were prepared from 1 μg total RNA input with a 200 bp median insert length. All libraries were quantified using a Qubit, pooled, and 100 bp paired-end reads were sequenced on two lanes of the HiSeq 2000 platform.

Raw RNA-seq data were screened to remove adapter sequences using Cutadapt v1.1 (Martin, 2011) with the following parameters: error-rate=0.1, times=1, overlap=5, and minimum-length=0. Adapter trimmed sequences were quality trimmed with Trimmomatic v0.22 (Bolger et al., 2014) using parameters (HEADCROP:0, LEADING:3, TRAILING:3, SLIDINGWINDOW:4:15, and MINLEN:40) to truncate reads for base quality <15 within 4 base windows and kept only reads ≥40 bases after trimming.

Reads were uniquely aligned the B73 RefGen_v2 maize genome assembly with GSNAP (Version 2013-07-20) using the following parameters: --orientation=FR -- batch=5 --suboptimal-levels=0 --novelsplicing=1 --localsplicedist=8000 --local-splice- penalty=0 --distant-splice-penalty=4 --quality-protocol=sanger --npaths=1 --quiet-if-excessive --max-mismatches=0.02 --nofails --format=sam --sam-multiple-primaries -- pairmax-rna=8000 --pairexpect=200 --pairdev=150 --nthreads=4. The --use-splicing argument was implemented to guide GSNAP by the ZmB73_5b Filtered Gene Set annotations but was permitted to discover novel splicing events. The argument --use-snps was called with an IIT file containing W22/B73 (Gault et al., 2017) as used to allow GSNAP to perform SNP tolerant alignments ensuring proper mapping of the w22 RNA-Seq reads to the B73 reference genome.

Read counts/gene were determined with the HTSeq-Count utility in the HTSeq package (Anders et al., 2015) (Ver 0.8.0). Non-redundant introns and genomic coordinates were identified from the ZmB73_5b annotation as described, except that U12-type introns were filtered to a non-redundant set of 372 introns (Gault et al., 2017) Percent spliced out (PSO) metrics for each intron were calculated from exon-exon junction spanning reads and intron reads for each unique 5’ and 3’ splice site (Gault et al., 2017; Katz et al., 2010). Fisher’s exact test was calculated for each intron from the summed exon-exon junction reads and intron reads across the four normal and four mutant libraries for each *rbm48* allele. Test statistics were correct to a false discovery rate ≤0.05 using the Benjamini-Hochberg method.

Differentially expressed transcripts were detected with the DESeq2 (Love et al., 2014) Bioconductor package. Transcripts were considered expressed if TPM was ≥1 in at least one genotype. Differential expression criteria were an adjusted p-value ≤0.05 and ≥2- fold change. GO term enrichment analysis used agriGO v2.0 with default parameters and the expressed transcripts as a customized reference (Du et al., 2010).

### Histology

Developing kernels from ears segregating for *rbm48* were harvested and fixed overnight at 4°C in FAA (3.7% formaldehyde, 5% glacial acetic acid, and 50% ethanol). Kernels were embedded in JB-4 plastic embedding media (Electron Microscopy Sciences) and sectioned at 4 μm thickness as described (Bai et al., 2016). Sections were stained in Schiff’s reagent, counter-stained with 1% aniline blue-black in 7% acetic acid, washed, dried, and mounted. Imaging was completed with a Zeiss Axiophot light microscope and an Amscope digital camera.

For β-glucuronidase (GUS) reporter expression analysis, hybrids of pBetl1:GUS or pVp1:GUS crossed by *rbm48*/+ heterozygous plants were self-pollinated (Hueros et al., 1999; Cao et al., 2007). Developing kernels were sampled from segregating ears, sectioned in the sagittal plane, and stained for GUS activity as previously described (Costa et al., 2003; Gutierrez-Marcos et al., 2006; Bai and Demason, 2008). At least 20 kernels for 3-4 biological replicates were observed, and 6-9 kernels were imaged to ensure that GUS reporter images were representative of the staining patterns. Images were captured with a Wild Heerbrugg dissecting microscope and an Amscope digital camera.

### Endosperm tissue culture

Endosperm callus cultures were initiated as described (Fouquet et al., 2011; Shannon, 1994). Briefly, ears segregating for *rbm48* from the A636 introgression were harvested from 6-18 DAP. Ears were surface sterilized in 70% ethanol and 20% bleach for 10 minutes and washed with sterile water. Individual endosperm tissues were dissected and placed on Murashige and Skoog media (pH 5.7) and supplemented with 3% sucrose, 4 ppm thiamine, 0.2% asparagine, and 2% phytagel. Growth of 400 individual endosperm tissues per genotype and harvest date was assayed after incubating in the dark for 35 d at 29°C. Cultures were imaged with a Nikon digital camera.

### Subcellular localization

RGH3, RGH3ΔUHM, U2AF1, and U2AF2 fusion protein constructs have been described previously (Gault et al., 2017; Fouquet et al., 2011). RBM48 fusion proteins were constructed with a similar strategy. Briefly, the full-length RBM48 ORF was amplified from B73 seedling cDNA. Primer pairs to enable Gateway cloning of N- or C-terminal fusions were used to amplify full-length (RBM48ΔFL), N-terminal deletion of the RRM domain (RBM48∆RRM), or C-terminal deletion (RBM48∆RS), see Supplementary Table 6 for primer sequences. PCR products were cloned into pDONR221 and recombined into Gateway destination vectors containing GFP, RFP, or split EYFP following the manufacturer’s protocol (Invitrogen) (Karimi et al., 2002; Karimi et al., 2007). For BiFC constructs, recombined expression cassettes, pSAT5-DEST or pSAT4-DEST, were digested with I-CeuI or I-SceI and ligated into pPZP-RCS2-bar binary vector. Binary vectors were transformed in *Agrobacterium tumefaciens* strains ABi and GV3101 (Wise et al., 2006).

*N. benthamiana* plants were grown for 5-8 weeks in a growth chamber 16/8 h and 26/22°C day/night for transient expression assays. *Agrobacterium* was infiltrated into stomata of the abaxial surface of leaves using a needleless syringe. Leaves were imaged 24-48 h after infiltration. For co-expression analysis, *Agrobacterium* strains with individual constructs were mixed in a 1:1 ratio prior to infiltration. Leaf sections were stained with DAPI prior to visualization of fusion protein expression (Kapila et al., 1997). Representative images of subcellular localization were obtained using a Carl Zeiss Axio Imager .Z2 confocal microscope (Cevik and Kazan, 2013).

### Protein pull-down assays

Coding sequences for maize U2AF1 (GRMZM2G177229) and U2AF2 (GRMZM2G093256) were amplified with primers containing Nco-I and Not-I restriction enzyme sites, cloned into a pCR4-TOPO vector (Invitrogen), then subcloned into pET-28b with standard restriction-ligation protocols. *E. coli* codon-optimized coding sequences of maize RBM48, RGH3, and ARMC7 were synthesized and cloned into pET-28b using NcoI and XhoI restriction sites (Genscript, Inc.). *Caldicellulosiruptor bescii* Cbes_2310 (WP_015908637.1), a predicted periplasmic sugar binding protein was cloned into pET-28b using NheI and XhoI restriction sites (Dam et al., 2011). A stop codon was included in the inserted *Cbes_2310* sequence to prevent translation of the C-terminal His-tag in the pET-28b vector. RBM48 and RGH3 were also cloned in pET-42a using NcoI and XhoI restriction sites to produce N-terminal tandem GST-His-tag fusion proteins (GenScript, Inc.). A stop codon was included in the inserted *RBM48* and *RGH3* sequences to prevent translation of the C-terminal His-tag in the pET-42a vector.

Recombinant expression conditions for all tagged versions of RGH3, U2AF1, U2AF2, and ARMC7 was initiated with a 1:50 dilution of overnight cultures in fresh LB media supplemented with 50 μg/mL kanamycin. Cultures were grown at 37°C with shaking at 225 rpm until OD_600_ = 0.45. Expression was induced by adding Isopropyl β-D-1-thiogalactopyranoside (IPTG) to a final concentration of 0.5 mM. Cultures were induced for 3 h at room temperature with shaking at 225 rpm. Bacterial cells were harvested by centrifugation at 2,500 x g for 15 min at 4°C. U2AF1, U2AF2, and ARMC7 cell pellets were stored at −80°C. RGH3 induction was improved by storing cell pellets overnight at 4°C with supernatant media. The supernatant was removed the following morning and the pellet stored at −80°C.

RBM48 recombinant proteins were expressed with a 1:20 dilution of an overnight grown culture into MagicMedia^TM^ *E. coli* Expression Medium (Invitrogen) supplemented with 50 μg/mL kanamycin. Cells were grown at 37°C with shaking at 225 rpm until OD_600_ = 0.6. The culture was then transferred to 18°C and grown for an additional 36 h at 225 rpm to allow for auto-induction of recombinant protein expression. After incubation at 18°C, the culture was centrifuged at 2,500 x g for 15 min at 4°C, and the pellet was stored at −80°C.

Cell pellets were resuspended in 2 mL of lysis buffer (50 mM Tris, 150 mM NaCl, 1X Halt^®^ Protease Inhibitor Cocktail (ThermoScientific), 1 mg/mL lysozyme, 10 μg/mL DNase I, 1%Triton X-100) and mixed with a rotisserie tube-rotator at room temperature for 10 min. The lysate was sonicated at 20 kHz for 30 s on/off five times, and subsequently centrifuged at 10,000 x g for 10 min at 4°C.

GST protein pull-down assays were based on Pierce^TM^ Glutathione Magnetic Agarose Beads instructions (ThermoScientific). Briefly, 400 μL bacterial lysate containing induced fusion protein was mixed with 100 μL equilibration buffer and incubated with 25 μL settled bead volume in a 1.5 mL microcentrifuge tube for 1 h at 4°C on a rotisserie tube-rotator. Beads were collected in a magnetic stand, the supernatant was discarded, and beads were washed twice with equilibration buffer. In the third wash, half of the slurry volume was transferred to a new microcentrifuge tube, and the GST fusion protein was eluted with 125 μL elution buffer to assay input protein for the pull-down.

The remaining protein-bound beads were separated on a magnetic stand, incubated with a mix of 200 μL of induced bacterial lysate containing His-tagged protein mixed with 50 μL of equilibration buffer. The lysate was incubated with the bound GST fusion protein for 1 h at 4°C on a rotisserie tube-rotator. The beads were collected and supernatant discarded. Beads were washed three times using 250 μL of equilibration buffer. Bound proteins were eluted with 125 μL elution buffer.

GST-His-RBM48, GST-His-RGH3, RGH3-His, U2AF1-His, U2AF2-His, ARMC7-His, and His-Cbes_2310 were detected by western blot using a monoclonal His-tag antibody (Cell Signaling, cat# 2365S). Protein fragments of RGH3-His were excised from the gels, subjected to trypsin digestion, and sequenced according to the procedure described (Shevchenko et al., 1996) at the Biomolecular and Proteomics Mass Spectrometry Facility at the University of California, San Diego.

## Accession numbers

All sequence data are available at the National Center for Biotechnology Information (NCBI) Short Read Archive (SRA). Maize transposon flanking sequence tags for identification of *rbm48-umu1* are in accession: SRX006230. Endosperm RNA-seq data are in accession: GSE118505.

## Acknowledgements

We thank John Baier, Jennifer Moses, J. Paige Gronevelt, Elizabeth Jankulovski, and Laurel Levine for technical assistance. This work was supported by National Science Foundation grants (MCB-1412218 to SL, WBB, AMS; IOS-1547787 to WBB; IOS-1623478 to AMS), the University of Florida HHMI Science for Life undergraduate research program, and the Vasil-Monsanto Endowment.

## Author contribution

A.M.S., S.L., W.B.B., F.B., and J.C. wrote the manuscript. A.M.S., F.B., G.S., and C.W.T. completed plant genetics and mutant mapping. F.B. characterized maize kernel, seedling, and cellular phenotypes. A.M.S., S.L., and G.F. completed protein sequence evolutionary analysis. W.B.B., R.D., F.B., A.M.S. designed and completed RNA-seq experiments. F.B., J.C., D.S., J.M., and A.E.S. completed RT-PCR experiments. A.M.S. and S.L. designed and supervised protein-protein interaction experiments. D.S. and F.M. completed co-localization and BiFC experiments. J.C., C.J.B., and F.M. developed recombinant expression constructs and methodology. J.C. completed protein pull-down experiments.

## Supplemental materials

Supplemental Figure 1. Molecular cloning of *rgh*-00F-061-05* to the *rbm48* locus (supports Figure 1).

Supplemental Figure 2. Phenotypes of *rbm48* alleles (supports Figure 1).

Supplemental Figure 3. Characterization of the *Rbm48* locus (supports Figures 1 and 2).

Supplemental Figure 4. RNA-seq read depth for experimentally validated genes (supports Figure 5).

Supplemental Figure 5. Overlap of RNA splicing defects in *rbm48* and *rgh3* mutant seedlings (supports Figure 5.)

Supplemental Figure 6. DEGs identified in *rbm48* mutant endosperm. (supports Figure 6).

Supplemental Figure 7. Endosperm cell differentiation defects in rbm48 at additional developmental stages (supports Figure 7).

Supplemental Table 1. Segregation of *rgh* kernel phenotypes in self-pollinations of *rbm48* heterozygotes (supports Figure 1).

Supplemental Table 2. Transmission of the *rbm48-umu1* allele in reciprocal crosses with normal inbred plants (supports Figure 1).

Supplemental Table 3. Read counts and PSO statistics (supports Figures 3-5).

Supplemental Table 4. Expressed transcripts (≥1 TPM in one genotype) with DeSeq2 adjusted p-values and DeSeq2 fold change ratios (supports Figure 6).

Supplemental Table 5. Enriched GO terms from differential gene expression analysis (supports Figure 6).

Supplemental Table 6. Primers used in this study (supports Methods section)

